## Supplementary Table 1 & Table 2 for "Eigenvector alignment: assessing functional network changes in amnestic mild cognitive impairment and Alzheimer’s disease"

### S1 Table

| | |
|---|---|
| 1 | Frontal Pole Right |
| 2 | Frontal Pole Left |
| 3 | Insular cortex Right |
| 4 | Insular cortex Left |
| 5 | Superior Frontal gyrus Right |
| 6 | Superior Frontal gyrus Left |
| 7 | Middle Frontal gyrus Right |
| 8 | Middle Frontal gyrus Left |
| 9 | Inferior Frontal gyrus, pars triangularis Right |
| 10 | Inferior Frontal gyrus, pars triangularis Left |
| 11 | Inferior Frontal gyrus, pars opercularis Right |
| 12 | Inferior Frontal gyrus, pars opercularis Left |
| 13 | Precentral gyrus Right |
| 14 | Precentral gyrus Left |
| 15 | Temporal Pole Right |
| 16 | Temporal Pole Left |
| 17 | Superior Temporal gyrus, anterior division Right |
| 18 | Superior Temporal gyrus, anterior division Left |
| 19 | Superior Temporal gyrus, posterior division Right |
| 20 | Superior Temporal gyrus, posterior division Left |
| 21 | Middle Temporal gyrus, anterior division Right |
| 22 | Middle Temporal gyrus, anterior division Left |
| 23 | Middle Temporal gyrus, posterior division Right |
| 24 | Middle Temporal gyrus, posterior division Left |
| 25 | Middle Temporal gyrus, temporooccipital part Right |
| 26 | Middle Temporal gyrus, temporooccipital part Left |
| 27 | Inferior Temporal gyrus, anterior division Right |
| 28 | Inferior Temporal gyrus, anterior division Left |
| 29 | Inferior Temporal gyrus, posterior division Right |
| 30 | Inferior Temporal gyrus, posterior division Left |
| 31 | Inferior Temporal gyrus, temporooccipital part Right |
| 32 | Inferior Temporal gyrus, temporooccipital part Left |
| 33 | Postcentral gyrus Right |
| 34 | Postcentral gyrus Left |
| 35 | Superior Parietal Lobule Right |
| 36 | Superior Parietal Lobule Left |
| 37 | Supramarginal gyrus, anterior division Right |
| 38 | Supramarginal gyrus, anterior division Left |
| 39 | Supramarginal gyrus, posterior division Right |
| 40 | Supramarginal gyrus, posterior division Left |
| 41 | Angular gyrus Right |
| 42 | Angular gyrus Left |
| 43 | Lateral Occipital cortex, superior division Right |
| 44 | Lateral Occipital cortex, superior division Left |
| 45 | Lateral Occipital cortex, inferior division Right |
| 46 | Lateral Occipital cortex, inferior division Left |

**Table S1.1.** A list of the ROIs (ID: 1 – 47) identified according to the CONN atlas.

| | |
|---|---|
| 47 | Intracalcarine cortex Right |
| 48 | Intracalcarine cortex Left |
| 49 | Frontal Medial cortex |
| 50 | Juxtapositional Lobule cortex -formerly Supplementary Motor cortex- Right |
| 51 | Juxtapositional Lobule cortex -formerly Supplementary Motor cortex- Left |
| 52 | Subcallosal cortex |
| 53 | Paracingulate gyrus Right |
| 54 | Paracingulate gyrus Left |
| 55 | Cingulate gyrus, anterior division |
| 56 | Cingulate gyrus, posterior division |
| 57 | Precuneus cortex |
| 58 | Cuneal cortex Right |
| 59 | Cuneal cortex Left |
| 60 | Frontal Orbital cortex Right |
| 61 | Frontal Orbital cortex Left |
| 62 | Parahippocampal gyrus, anterior division Right |
| 63 | Parahippocampal gyrus, anterior division Left |
| 64 | Parahippocampal gyrus, posterior division Right |
| 65 | Parahippocampal gyrus, posterior division Left |
| 66 | Lingual gyrus Right |
| 67 | Lingual gyrus Left |
| 68 | Temporal Fusiform cortex, anterior division Right |
| 69 | Temporal Fusiform cortex, anterior division Left |
| 70 | Temporal Fusiform cortex, posterior division Right |
| 71 | Temporal Fusiform cortex, posterior division Left |
| 72 | Temporal Occipital Fusiform cortex Right |
| 73 | Temporal Occipital Fusiform cortex Left |
| 74 | Occipital Fusiform gyrus Right |
| 75 | Occipital Fusiform gyrus Left |
| 76 | Frontal Operculum cortex Right |
| 77 | Frontal Operculum cortex Left |
| 78 | Central Opercular cortex Right |
| 79 | Central Opercular cortex Left |
| 80 | Parietal Operculum cortex Right |
| 81 | Parietal Operculum cortex Left |
| 82 | Planum Polare Right |
| 83 | Planum Polare Left |
| 84 | Heschl's gyrus Right |
| 85 | Heschl's gyrus Left |
| 86 | Planum Temporale Right |
| 87 | Planum Temporale Left |
| 88 | Supracalcarine cortex Right |
| 89 | Supracalcarine cortex Left |

**Table S1.2.** A list of the ROIs (ID: 47 – 89) identified according to the CONN atlas.

| | |
|---|---|
| 90 | Occipital Pole Right |
| 91 | Occipital Pole Left |
| 92 | Thalamus Right |
| 93 | Thalamus Left |
| 94 | Caudate Right |
| 95 | Caudate Left |
| 96 | Putamen Right |
| 97 | Putamen Left |
| 98 | Pallidum Right |
| 99 | Pallidum Left |
| 100 | Hippocampus Right |
| 101 | Hippocampus Left |
| 102 | Amygdala Right |
| 103 | Amygdala Left |
| 104 | Accumbens Right |
| 105 | Accumbens Left |
| 106 | Brainstem |
| 107 | Cerebellum Crus1 Left |
| 108 | Cerebellum Crus1 Right |
| 109 | Cerebellum Crus2 Left |
| 110 | Cerebellum Crus2 Right |
| 111 | Cerebellum 3 Left |
| 112 | Cerebellum 3 Right |
| 113 | Cerebellum 4 5 Left |
| 114 | Cerebellum 4 5 Right |
| 115 | Cerebellum 6 Left |
| 116 | Cerebellum 6 Right |
| 117 | Cerebellum 7b Left |
| 118 | Cerebellum 7b Right |
| 119 | Cerebellum 8 Left |
| 120 | Cerebellum 8 Right |
| 121 | Cerebellum 9 Left |
| 122 | Cerebellum 9 Right |
| 123 | Cerebellum 10 Left |
| 124 | Cerebellum 10 Right |
| 125 | Vermis 1 2 |
| 126 | Vermis 3 |
| 127 | Vermis 4 5 |
| 128 | Vermis 6 |
| 129 | Vermis 7 |
| 130 | Vermis 8 |
| 131 | Vermis 9 |
| 132 | Vermis 10 |

**Table S1.3.** A list of the ROIs (ID: 90 – 132) identified according to the CONN atlas.

#### S2 Table

| ID | 1 | 2 | 3 | ID | 1 | 2 | 3 | ID | 1 | 2 | 3 |
|---|---|---|---|---|---|---|---|---|---|---|---|
| 1 | 8 | 4 | 10 | 47 | 1 | 4 | 3 | 93 | 4 | 5 | 6 |
| 2 | 5 | 2 | 2 | 48 | 1 | 4 | 6 | 94 | 2 | 2 | 0 |
| 3 | 6 | 4 | 6 | 49 | 4 | 4 | 4 | 95 | 4 | 10 | 1 |
| 4 | 13 | 3 | 2 | 50 | 9 | 5 | 7 | 96 | 2 | 4 | 3 |
| 5 | 6 | 3 | 13 | 51 | 11 | 4 | 4 | 97 | 2 | 3 | 3 |
| 6 | 5 | 0 | 7 | 52 | 6 | 3 | 2 | 98 | 3 | 1 | 0 |
| 7 | 2 | 4 | 3 | 53 | 8 | 5 | 2 | 99 | 5 | 21 | 9 |
| 8 | 5 | 5 | 3 | 54 | 13 | 9 | 2 | 100 | 2 | 2 | 8 |
| 9 | 5 | 3 | 1 | 55 | 0 | 3 | 5 | 101 | 2 | 7 | 13 |
| 10 | 4 | 2 | 7 | 56 | 1 | 4 | 1 | 102 | 2 | 8 | 8 |
| 11 | 4 | 4 | 5 | 57 | 1 | 2 | 4 | 103 | 5 | 1 | 3 |
| 12 | 4 | 6 | 11 | 58 | 2 | 9 | 6 | 104 | 4 | 19 | 4 |
| 13 | 4 | 4 | 4 | 59 | 2 | 6 | 1 | 105 | 7 | 16 | 4 |
| 14 | 2 | 3 | 2 | 60 | 11 | 7 | 4 | 106 | 5 | 6 | 7 |
| 15 | 4 | 4 | 7 | 61 | 9 | 3 | 7 | 107 | 0 | 9 | 7 |
| 16 | 4 | 1 | 7 | 62 | 7 | 4 | 5 | 108 | 4 | 3 | 2 |
| 17 | 2 | 1 | 3 | 63 | 16 | 4 | 8 | 109 | 7 | 1 | 6 |
| 18 | 3 | 3 | 3 | 64 | 2 | 21 | 11 | 110 | 2 | 11 | 19 |
| 19 | 9 | 3 | 3 | 65 | 1 | 4 | 3 | 111 | 2 | 1 | 2 |
| 20 | 4 | 9 | 4 | 66 | 7 | 9 | 3 | 112 | 6 | 1 | 3 |
| 21 | 6 | 2 | 6 | 67 | 4 | 3 | 6 | 113 | 8 | 3 | 2 |
| 22 | 7 | 2 | 8 | 68 | 8 | 3 | 12 | 114 | 2 | 7 | 4 |
| 23 | 5 | 19 | 11 | 69 | 7 | 3 | 6 | 115 | 6 | 6 | 13 |
| 24 | 2 | 12 | 15 | 70 | 6 | 9 | 16 | 116 | 4 | 8 | 6 |
| 25 | 1 | 4 | 0 | 71 | 4 | 7 | 10 | 117 | 1 | 4 | 1 |
| 26 | 2 | 1 | 2 | 72 | 6 | 7 | 6 | 118 | 3 | 6 | 4 |
| 27 | 5 | 3 | 6 | 73 | 4 | 6 | 10 | 119 | 3 | 6 | 7 |
| 28 | 3 | 7 | 3 | 74 | 1 | 4 | 9 | 120 | 2 | 4 | 5 |
| 29 | 7 | 3 | 15 | 75 | 1 | 6 | 5 | 121 | 3 | 3 | 9 |
| 30 | 5 | 13 | 22 | 76 | 3 | 6 | 5 | 122 | 4 | 1 | 8 |
| 31 | 3 | 7 | 19 | 77 | 3 | 4 | 7 | 123 | 10 | 11 | 8 |
| 32 | 5 | 9 | 8 | 78 | 3 | 4 | 5 | 124 | 2 | 4 | 6 |
| 33 | 4 | 3 | 6 | 79 | 5 | 10 | 12 | 125 | 3 | 1 | 6 |
| 34 | 6 | 3 | 3 | 80 | 0 | 7 | 3 | 126 | 1 | 1 | 2 |
| 35 | 5 | 1 | 1 | 81 | 3 | 4 | 10 | 127 | 5 | 2 | 1 |
| 36 | 5 | 2 | 2 | 82 | 15 | 4 | 14 | 128 | 8 | 9 | 13 |
| 37 | 2 | 4 | 1 | 83 | 12 | 10 | 15 | 129 | 3 | 7 | 6 |
| 38 | 1 | 3 | 4 | 84 | 9 | 4 | 9 | 130 | 5 | 7 | 3 |
| 39 | 0 | 5 | 2 | 85 | 21 | 4 | 8 | 131 | 3 | 3 | 3 |
| 40 | 2 | 1 | 6 | 86 | 8 | 7 | 9 | 132 | 0 | 9 | 1 |
| 41 | 5 | 3 | 2 | 87 | 6 | 3 | 8 | | | | |
| 42 | 2 | 5 | 4 | 88 | 2 | 7 | 6 | | | | |
| 43 | 2 | 1 | 2 | 89 | 9 | 5 | 5 | | | | |
| 44 | 12 | 12 | 4 | 90 | 3 | 5 | 4 | | | | |
| 45 | 2 | 3 | 7 | 91 | 2 | 8 | 16 | | | | |
| 46 | 4 | 5 | 5 | 92 | 3 | 2 | 3 | | | | |

**Table S2.1.** Eigenvector alignment where the number of significant changes in eigenvector alignment for each ROI with respect to other ROI are detailed for the comparisons (1) AD versus HC, (2) aMCI versus HC, (3) AD versus aMCI.
